# Using kinetic modelling to infer adaptations in *Saccharomyces cerevisiae* carbohydrate storage metabolism to feast famine regimes

**DOI:** 10.1101/2022.06.28.497175

**Authors:** David Lao-Martil, Koen J.A. Verhagen, Ana H. Valdeira Caetano, Ilse H. Pardijs, Natal A.W. van Riel, S. Aljoscha Wahl

## Abstract

Microbial metabolism is strongly dependent on the environmental conditions. While these can be well controlled under laboratory conditions, large-scale bioreactors are characterized by inhomogeneities and consequently dynamic conditions for the organisms. How *Saccharomyces cerevisiae* responds to frequent perturbations in industrial bioreactors is still not understood mechanistically. To study the adjustments to prolonged dynamic conditions, experiments under a feast/famine regime were performed and analysed using modelling approaches. Multiple types of data were integrated; including quantitative metabolomics, ^13^C incorporation and flux quantification. Kinetic metabolic modelling was applied to unravel the relevant intracellular metabolic response mechanisms. An existing model of yeast central carbon metabolism was extended, and different subsets of enzymatic kinetic constants were estimated. A novel parameter estimation pipeline based on combinatorial enzyme selection, supplemented by regularization, was developed to identify and predict the minimum enzyme and parameter adjustments from steady-state to feast famine conditions. This approach predicted proteomic changes in hexose transport and phosphorylation reactions, which was additionally confirmed by proteome measurements. Nevertheless, the modelling also hints to a yet unknown kinetic or regulation phenomenon. Some intracellular fluxes could not be reproduced by mechanistic rate laws, including hexose transport and intracellular trehalase activity during feast famine cycles.

**Author summary:** Kinetic metabolic models are used to understand how biological systems deal with dynamic perturbations in their environment. A well-known case of their application is the microorganism *Saccharomyces cerevisiae*, which was domesticated by mankind thousands of years ago, and is used to produce a wide range of products such as bread, beverages, and biofuels. When cultured in industrial-scale bioreactors, this cell factory is impacted by environmental perturbations which can challenge the bioprocess performance. The feast famine regime has been proposed as an experimental setup to downscale these industrial perturbations. Intracellularly, these perturbations impact central carbon metabolism, including carbon storage. Even though kinetic metabolic models have been developed to study the effect of extracellular perturbations, they have not explored the feast famine regime and its implications on carbon metabolism. We developed a model identification tool and used it to expand the existing models to represent carbon metabolism under feast famine regime. We used computer simulations to point at adaptations in yeast metabolism and locations in the model where our understanding is not entirely accurate. We found that combining multiple types of data, despite challenging, can be very beneficial by providing a comprehensive and realistic representation of the cell.

## Introduction

*Saccharomyces cerevisiae*, also commonly known as baker’s yeast, has been used by mankind for thousands of years for the production of relevant beverages, foods and chemicals. However, despite its extensive usage in industry [1–3], scaling especially new *S. cerevisiae* production processes to industrial scale poses several, interesting and fundamental challenges. Source of most challenges is spatial inhomogeneities due to mixing limitations in large-scale bioreactors leading to gradients throughout the reactor. A cell dispersed in the reactor is therefore exposed to rapid changes in its extracellular environment, which in turn will impact intracellular metabolic regulation [4,5]. Similarly, the natural habitat will commonly have oscillations resp. perturbations of environmental conditions like temperature, pH and substrate availabilities.

Although dynamic conditions are encountered for industrial applications as well as environmental habitats, many physiological studies on yeast are performed under (pseudo-) steady-state conditions. Clearly, with the vast available reference data, reliable measurements, and reproducibility, steady-state (SS) experiments are very useful in the quantification of intracellular fluxes. However, for the identification of in vivo kinetic parameters, dynamic metabolic experiments are required [6]. To bridge this gap, dynamic perturbation experiments can be performed, and many studies have focused on elucidating the metabolic response from single pulse (SP) experiments [6–12].

While this SP approach is very useful for the identification of kinetic parameters of networks adapted to the pre-perturbation limited steady-state, it cannot describe adaptations that may occur upon the subsequent perturbations observed under industrial conditions [5]. To emulate such an environment, a system of periodic perturbations, known as a feast/famine (FF) regime, was developed. The FF regime produces repetitive substrate concentration gradients in time, which allow for accurate and reproducible sampling of the intracellular metabolism (Fig 1) [13, 14]. Suarez-Mendez et al. (2014) used this FF setup to monitor the *in vivo* metabolic activity during FF cycles of 400 s. At this timescale it is assumed that the metabolic response within one cycle is mainly governed by metabolic interactions, as enzyme concentrations will remain basically constant during these 400 s [15]. In a FF cycle, feed was provided block-wise i.e. 20 s feeding, followed by 380 s of no feed (Fig 1).

**Fig 1.**
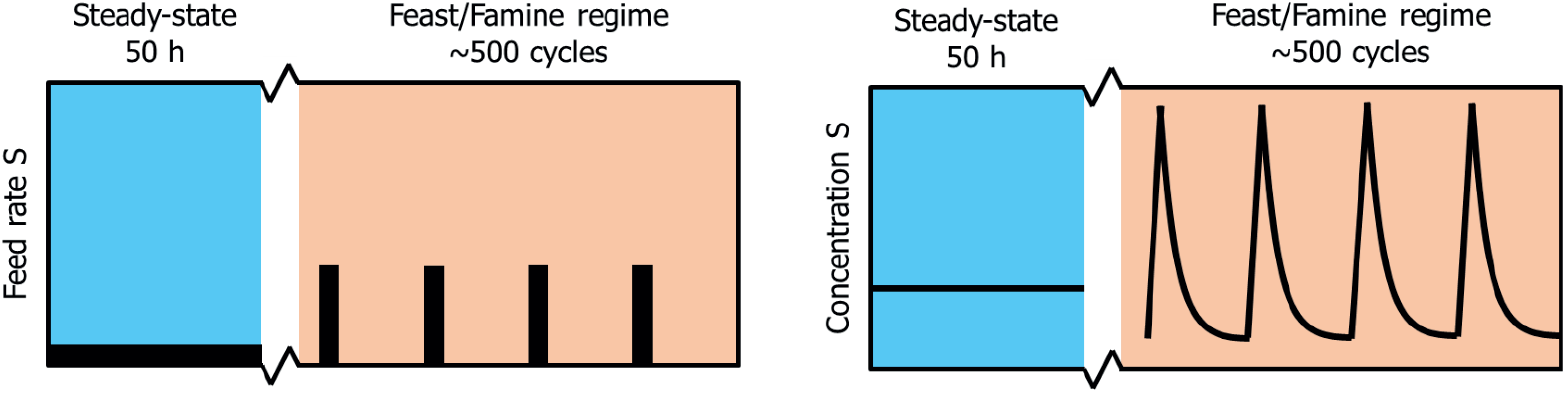
Profile of the experimental feeding regime. After a chemostat phase (reference steady-state) of 50 h, a block-wise feed is applied in a 400 s cycle at the same average substrate supply and dilution rate for another 50 h (adapted from [13]).

Under such FF conditions, *S. cerevisiae* cultures show different metabolic phenotypes compared to SP or SS cultures [13]. After a pulse in FF, an increase rather than a decrease in ATP, no ethanol production, and no accumulation of glycolytic metabolites was observed. These differences in metabolic response suggest a proteomic adaptation, induced by the prior dynamic growth conditions [16–18]. Especially, translational regulation can lead to condition specific proteome compositions [19]. In fact, distinct proteome compositions have been observed under changing glucose availability conditions [20], both for sugar transporters and intracellular enzymes [21, 22], and distinct isoenzymes have different kinetic properties that can include glucose sensitivity as well [23]. However, the mechanisms behind this adaptation are not well understood.

For the identification of kinetic parameters and putative regulation mechanisms, quantitative data from different approaches will be required. Especially, to generate comprehensive models, in addition to intracellular concentrations, carbon tracing is required to identify bidirectional, cyclic or parallel reactions [24, 25]. Work with 13C-labelling indicated that storage metabolism is a major metabolic sink upon changes in the glycolytic flux, with in average 15% of the carbon flux being diverted through the glycogen and trehalose cycles [26]. This diversion of flux is in accordance with earlier studies into the importance of the trehalose cycle under SP conditions [9].

Understanding metabolism, especially under dynamic conditions requires integration of stoichiometry and enzyme kinetics. Such kinetic metabolic models have been developed for *S. cerevisiae* using either *in vitro* and *in vivo* parameters [22, 27], additional allosteric regulation [28] or subpopulation dynamics [9]. Nonetheless, the conditioning of the cells was mostly at steady-state [29], leading to putative mismatches when applied to large-scale, dynamic cultivation conditions.

Under dynamic conditions, further pathways have been described to play a regulatory role – glycogen and trehalose metabolism. For both, the reaction stoichiometry is known, and *in vitro* parameters have been broadly studied [30–33], but no *in vivo* based parameter values have been derived.

Especially for cyclic pathways like the trehalose cycle, quantifying *in vivo* parameters can be challenging as both in- and outfluxes influence the concentration change and no in- or outflux is directly observable. This correlation, plus the fact that the networks are getting larger, leads to a danger posed by local minima and ill-conditioning, and consequently sloppy parameter estimates [34]. To overcome this challenge and identify a minimal set of necessary changes in kinetic constants, the divide and conquer approach has been developed. Here, a decomposition of the global estimation problem into independent subproblems [35] is used. Furthermore, to consider the already known parameter values for the enzymes under study [22,29], L1 or Tikhonov regularization can favour a given parameter set, as long as experimental data is properly reproduced [36–38].

Here, we specifically studied the impact of proteome adaptation to substrate perturbations on the changed metabolic response. To this end, we have expanded upon existing state-of-the-art kinetic models, integrating both metabolomics and fluxomics as well as ^13^C enrichment data, to evaluate which proteome changes are most relevant to explain the experimentally observed change in metabolic response.

## Results and Discussion

### Continuous and feast famine grown cells demonstrate different enzymatic levels and metabolic responses – experimental observations

As mentioned before, cells exposed to a block-wise feeding (feast/famine conditions), showed a remarkable different response compared to glucose-limited cells from chemostat conditions. The proteome during both conditions was measured and subsequently analysed on their composition [39]. Major changes were observed within the glycolytic and transporter enzymes, specifically in the expression levels of hexose transporters (HXT), hexokinase (HXK) and glyceraldehyde dehydrogenase (TDH) (Fig 2). The expression of hexose transporters, especially HXT7, has been shown to correlate with the (maximal) substrate uptake rate [40]. The observed decrease in protein concentration can be interpreted as an adaptation to limit rapid influx of glucose upon glucose pulse. In contrast, glucokinase (GLK) is slightly upregulated. HXK is highly regulation through inhibition by T6P, however GLK is not inhibited by trehalose-6-P (T6P) up to a level of 5 mM [41]. As such the lower concentration of HXK in combination with the upregulation of GLK will likely result in an adaptation in the regulation of the glycolytic flux. The downregulation of upper glycolysis (HXK), in combination with the upregulation of lower glycolysis (TDH) may additionally allow for improvement of flux capacity through glycolysis upon glucose influx [9].

**Fig 2.**
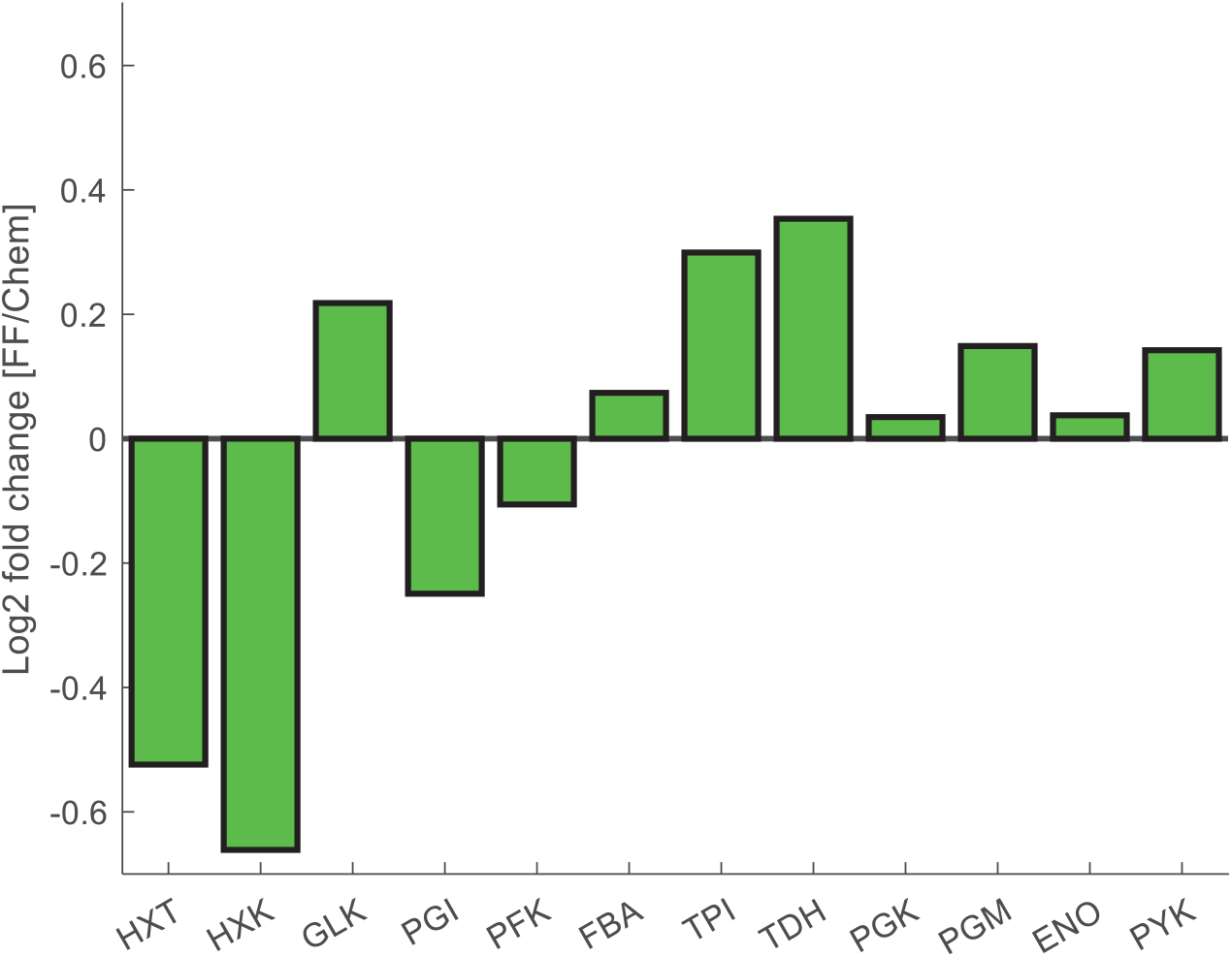
The protein concentration log2 fold change from chemostat to feast/famine conditions of selected glycolytic and transporter proteins.

[42] observed that long-term adaptation of the proteome composition had a major influence on the adjustment of the metabolic response of the cell. To assess whether the observed metabolic response can indeed be explained by the measured proteome changes, combinatorial enzyme selection and regularization was used to identify key parameter adaptations. Next to this, the effect of individual iso-enzymes was considered and evaluated as a factor influencing the observed metabolic response. Furthermore, the kinetics and implementation of the storage metabolism were evaluated.

### Carbon storage physiology differs between continuous and feast famine limitation

A kinetic model of yeast glycolysis was previously developed to fit various SS and SP data sets [29]. For the FF conditions, the model was extended with regard to trehalose metabolism in different compartments and glycogen synthesis and degradation (Fig 3).

**Fig 3.**
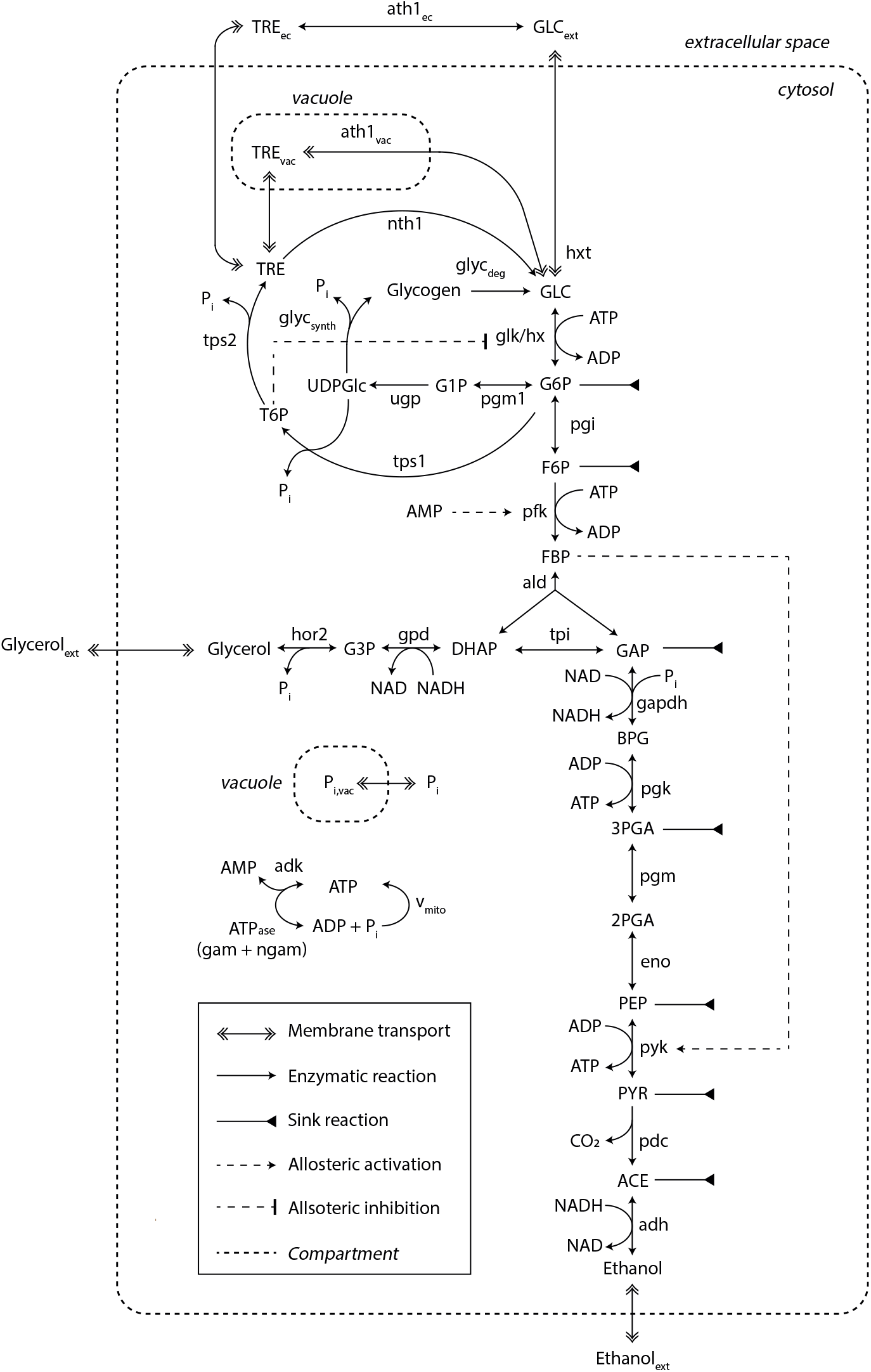
Kinetic metabolic model with a detailed description of trehalose and glycogen metabolism. Sink reactions account for fluxes towards the TCA, PPP and biomass synthesis. This model was adapted from [29]. The diagram style was adapted from [9]. For a detailed view on model mass balances and reaction kinetics, see supplementary S1 Appendix and S2 Appendix.

The model simulations could reproduce most of the experimental data properly, after several kinetic constants from HXT and HXK were estimated (this is explained in detail in the next section), but with a few exceptions (Fig 4A). For instance, glucose 6-phosphate (G6P) and fructose 6-phosphate (F6P) simulated concentrations were smaller, which was already documented in the original model and attributed to underdetermined phosphofructokinase (PFK) reaction kinetics [29]. Other metabolites also deviated, such as fructose bis-phosphate (FBP), glucose 1-phosphate (G1P) or trehalose 6-phosphate (T6P). This could be explained by affinity constants undergoing changes during FF cycles, here unaccounted for. Furthermore, reaction rates, estimated from ^13^C enrichment data, were in close agreement (Fig 4B), except for the maximum simulated rate for HXT with a simulated maximum lower than observed experimentally.

**Fig 4.**
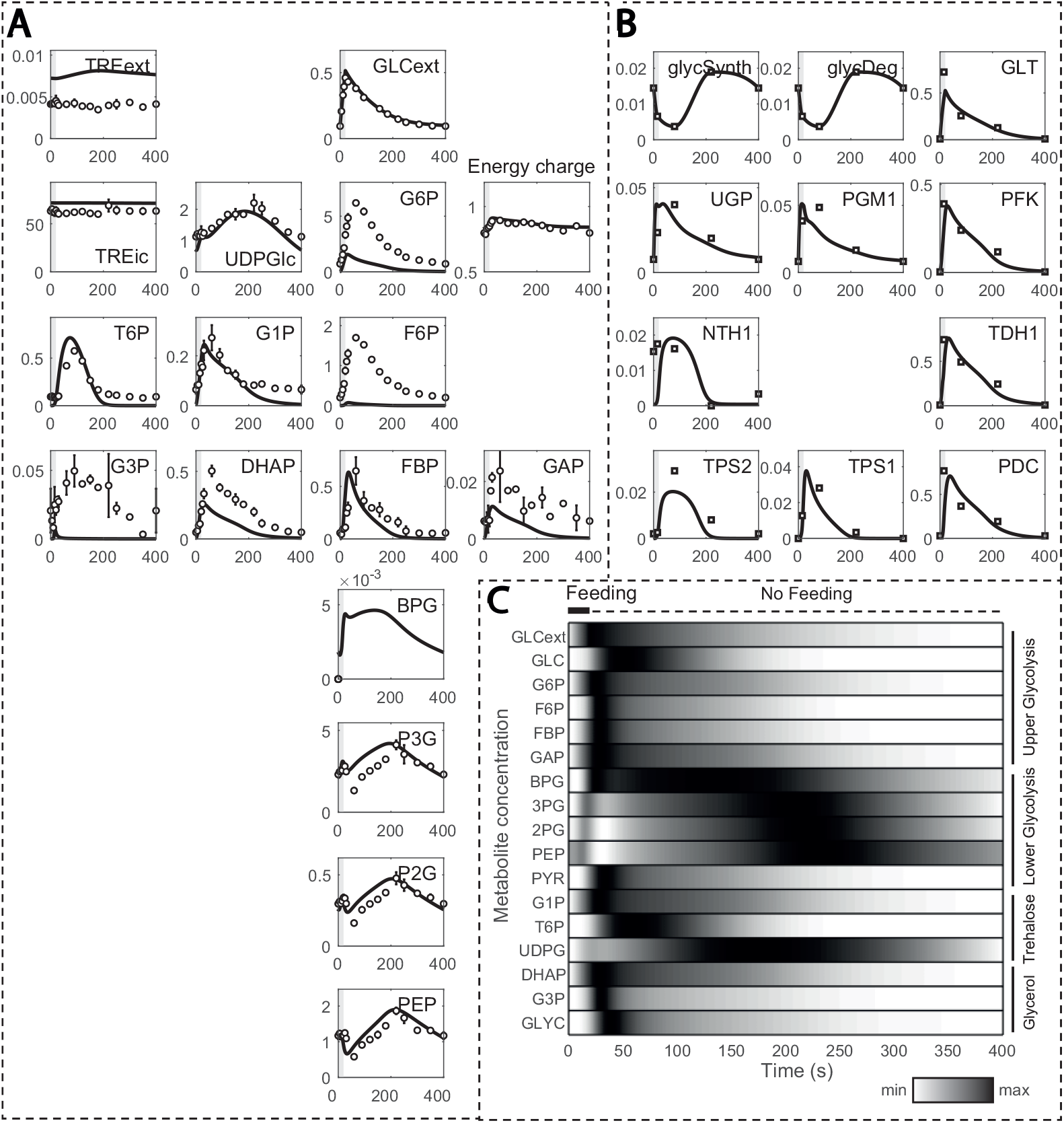
Model simulations in comparison to experimental observations. **(A) Metabolite concentrations and (B) reaction rates estimated from 13C enrichment data over one cycle (400 s)**. Metabolite concentrations and reaction rates are displayed in the y-axis (in mM and mM s^-1^, respectively) and time in the x-axis. **(C) Normalized metabolite concentrations during one feeding cycle**. Darker colours indicate values closer to the maximum, while brighter ones to the minimum.

In the FF simulations, the increase in residual glucose during feeding resulted in transient changes in glycolytic metabolites, which returned to the initial value by the end of the cycle (Fig 4C). Upper glycolysis metabolites reached their maximum concentration within 50 seconds, except glucose, due to recirculation via storage metabolism. Glycerol branch and storage kinetics followed a similar but with delay. Due to the slower reaction rate for enzyme GAPDH [43], the entry in lower glycolysis was delayed and BPG reaches its maximum at about 130 seconds. Nonetheless, the increase in FBP activated pyruvate kinase (PYK), reducing concentrations of 3-phosphoglycerate (P3G), 2-phosphoglycerate (P2G) and phosphoenolpyruvate (PEP). As FBP decreased, its activation dissipated, and these lower glycolysis metabolites reached maximum concentrations in about 230 to 250 seconds of cycle. This trend is in agreement with the known allosteric regulation of FBP on PTK [12].

Trehalose metabolic dynamics were different FF and SP. During FF, the maximum flux towards production of trehalose was less than 10% of the HXK reaction rate, in comparison to the 30% observed in SP [9], and part was secreted to the extracellular space, what is commonly regarded as stress protection [44, 45]. Nonetheless, glycogen took up a greater portion during the feast famine regime, implying that carbon storage is predominant over the stress response by the trehalose cycle and suggesting that the cell is indeed adapted to the FF setup.

Small changes in protein expression occur between cells in a population. One way to examine the possible resulting phenotypes of the network upon perturbation is by means of ensemble modelling [46, 47]. To test the robustness of the model, 10000 simulations were performed with random parameter values deviating within a range of 10% of the model parameter set. The concentration and reaction rate profiles were very consistent (see supplementary S1 Fig andS2 Fig, respectively), especially for reaction rates, where the relative deviation between fluxes was very small. This suggested that the model dynamics are consistent within the parameter range tested.

### Glucose transport and phosphorylation identified as key adaptations from combinatorial parameter estimation

Glycolytic enzymes expression changed from chemostat to feast/famine conditions, most notably for HXT and HXK (Fig 2). As each iso-enzyme has specific kinetic properties, the catalytic (*K*_*cat*_) and Michaelis-Menten (*K*_*M*_) constants also differ [22]. To identify changes in kinetic parameters, parameter estimation based on the FF datasets was performed and interpreted in light of the measured proteome changes. However, estimating all kinetic parameters simultaneously can lead to multiple local minima and ill-conditioning [34]. To bypass this problem and identify which are the key parameters that change between the cells adapted to continuous and feast famine conditions respectively, we adapted the scale and setup of the parameter estimation problem. Two stages were applied (Fig 5): (1) Parameters were estimated for multiple combination of enzymes in parallel assays, to isolate which enzymes were key to reproduce the data properly, and (2) Regularization was implemented, i.e., parameters were re-estimated for the best selection of enzymes found and with a penalty for deviation from the reference parameter set.

**Fig 5.**
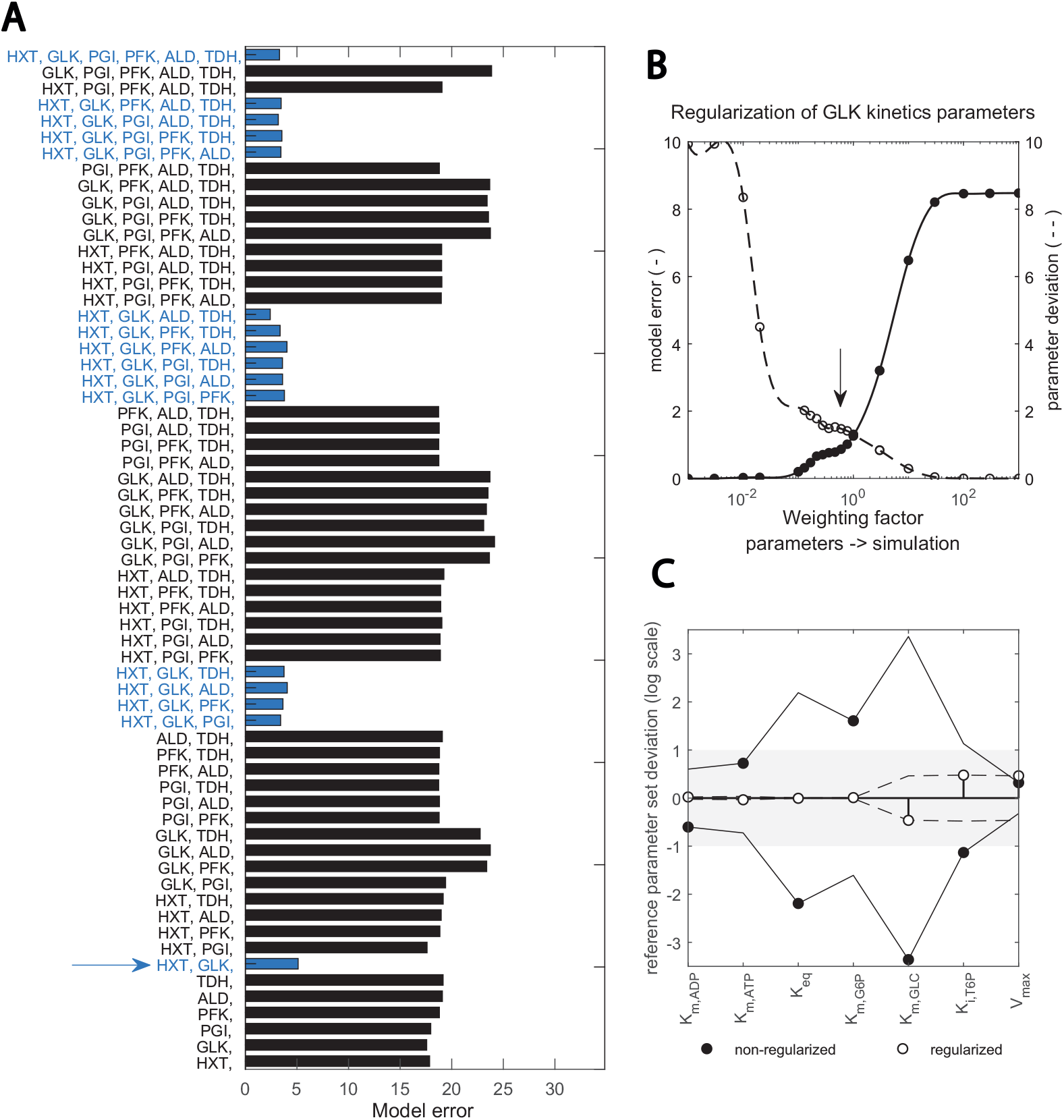
Two-step, scaled optimization approach results: **(A) Fitting the data with different subsets of enzyme parameters**. Bars show the error between simulation and best fit with the respective combination (upper x-axis). Blue bars highlight the combinations containing the two enzymes HXT and GLK. Red lines show the number of enzymes. **(B) Implementation of a regularization factor on the estimation of GLK kinetic parameters**. The dashed and continuous line show model and parameter error, respectively. The arrow indicates the chosen regularization factor. Parameters are regularized such that the data is still well reproduced (see supplementary S3 Fig and S4 Fig). **(C) Change in key GLK parameters identified upon regularization**. The deviation between the estimated parameter and the initial value taken form (bioRxiv2022) is shown in the y-axis (in logarithmic scale). Black and empty circles show the estimates prior and post regularization, when parameter dependencies are minimized. The x-axis shows specific parameters.

In step 1 (Fig 5A), good fits were achieved when HXT, HXK/GLK and the trehalose cycle were included in the parameter estimation, suggesting that these are the relevant enzymes undergoing changes. In step 2, an optimal fit between model error and parameter deviation was achieved by adding a regularization factor, shown in (Fig 5B) for GLK kinetics. This overcame dependencies between kinetic constants of the reaction and pointed to *K*_*m,GLC*_, *K*_*i,T*6*P*_ and *V*_*max*_ being the key parameter alterations with respect to the reference parameter set from [29] (Fig 5C). Compared with other toolboxes available to perform parameter estimation in complex kinetic metabolic models [22, 34, 36, 48, 49], this pipeline incorporates regularization for known *in vitro* parameter values into a combinatorial enzyme selection approach, with the added value that intracellular flux data is used. Flux data has been available only for a decade and few works have used it for yeast kinetic model development and validation [9, 29].

As a result of this pipeline, some kinetic constant changes were suggested for the glucose transport and phosphorylation reactions Table 1. For HXT, the maximal reaction rate (*V*_*max*_) decreased from 8.13 to 1.7 mM s^-1^, which is actually close to the value in other published models [27, 28] and consistent with the experimental HXT concentration decrease (Fig 2). Changes in HXT isoenzymes proportions could also lead to changes in affinity; *In-vitro K*_*M*_ measurements have shown a high variability for the HXT1-7 subunits [21, 50]. Nevertheless, the shift in isoenzymes here did not lead to drastic changes in affinity, the *K*_*M*_ did only change from 1.01 to 0.90 mM.

**Table 1.**
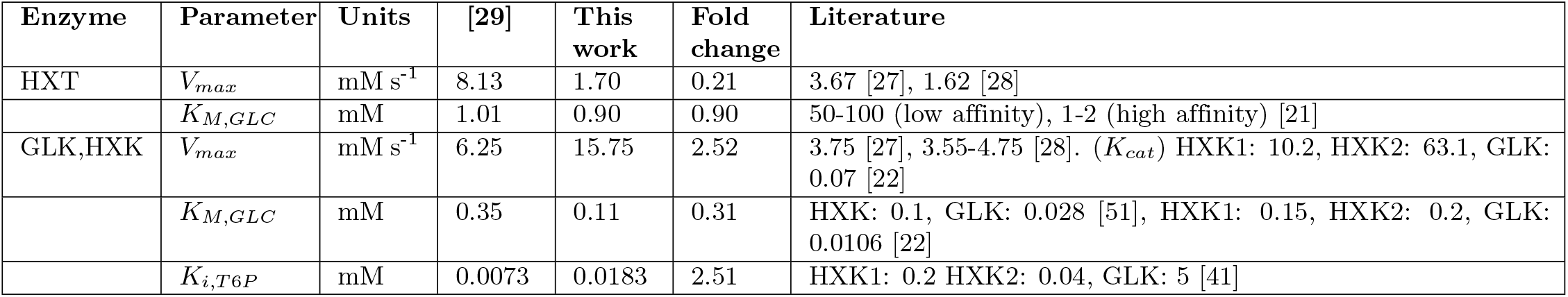
Table 1 Several parameters explain the adaptations in HXT and HXK isoenzymes. Changes in parameters in HXT and HXK kinetics that allow the Y3M1 model to fit the data. For the other parameters in these reactions, changes were below 5%.

For the hexose phosphorylation reaction, several kinetic constants changed. Km,glucose decreased from 0.35 to 0.11 mM, in line with the experimental increase in GLK/HXK ratio (Fig 2), given that the affinity constant for glucose was found lower in GLK than HXK both in vitro and *in vivo*-like conditions [22, 51]. Furthermore, the T6P inhibition constant (*K*_*i*_) increased from 0.0073 to 0.0183 and *K*_*i,T*6*P*_ was also found to change between the two isoenzymes [41]. Next to these parameters, *V*_*max*_ increased from 6.25 to 15.75 mM s^-1^, which is higher than values reported before in yeast glycolytic models [27, 28]. Since the *k*_*cat*_ for GLK is much lower than for HXK [22] such increase in *V*_*max*_ was not expected.

Outside glycolysis, changes were required in the trehalose cycle and ATP maintenance reaction. Note, that the original model did not include glycogen metabolism, nor compartmentation of the trehalose cycle reactions. The missing reactions were added in this work, and trehalose cycle parameters re-estimated to account for the effect of the previously lumped reactions. Finally, the ATPase maximum reaction rate decreased to fit the adenosine nucleotide concentrations (ATP + ADP + AMP). This might be related to the fact that the initial model simulated the response to a GP of 20 g L^-1^ of glucose [29], which is seen by the cell as a stress condition [9]. The final parameter set can be found in the supplementary S3 Appendix.

### Glucose sensing influences hexose transporter kinetics during feast-famine cycles

Glucose uptake has been widely modelled as an equilibrium-driven passive transport reaction [22, 27], where its kinetics are determined by isoenzyme-specific *V*_*max*_ and *K*_*M*_ parameters [21, 51, 52]. Here, we have found that these kinetics alone cannot explain the experimental data, for which a glucose sensing mechanism [23] needs to be active.

By sampling the parameter space and using passive transport reaction kinetics, we found that no combination of parameters could fit the data (Fig 6). Especially, none of the generated models could reproduce a net uptake reaction of almost zero at the end of the cycle (400 seconds) and reach the value of 0.72 mM s^-1^ at 20 seconds, when the uptake rate reaches its maximum (Fig 6A). Since throughout the entire FF cycle the residual and maximum glucose concentration are 0.1 and 0.45 g L^-1^, respectively, adjusting parameters to lower the effect of transmembrane glucose gradient for one also reduces for the other. Interestingly, the only way to reproduce the experimental uptake (Fig 6B) was by including a minimum glucose concentration term in HXT kinetics (Csmin in equation 2). This term acted as a threshold value from above which glucose import occurs.

**Fig 6.**
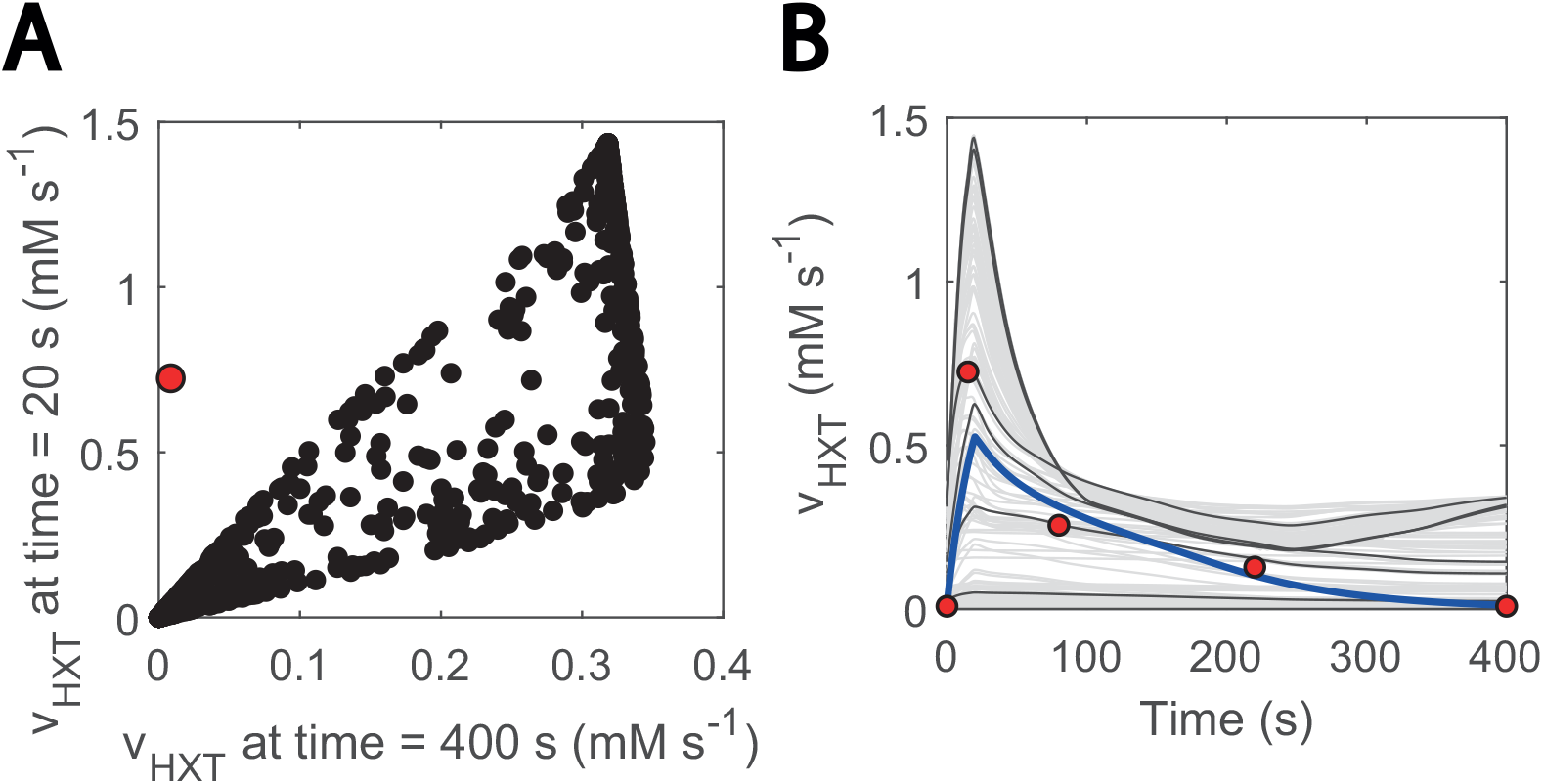
Glucose sensing is needed to explain HXT kinetics. **(A) Glucose uptake rate at 20 seconds vs at 400 seconds.** 20 seconds is the approximate point for the maximum reaction rate. Black data shows simulations generated with randomly generate parameter samples when no threshold value is considered. 1000 samples were run, within 3 orders of magnitude above and below the estimated parameters. Parameters were randomized for HXT kinetics and external glucose concentration was fit to the experimental data. **(B) Visualization of hexose transport rate during the cycle for the abovementioned models**. The blue line corresponds to the simulation with the model considers glucose sensing. The grey and black coloured simulations are the ones with the generated models. Only 200 are displayed and some simulations are highlighted in black to ease visualization. The red dots point at the experimental data points. The individual effect of HXT kinetic parameters (*V*_*max*_, *K*_*M*_) can be found in supplementary S5 Fig.

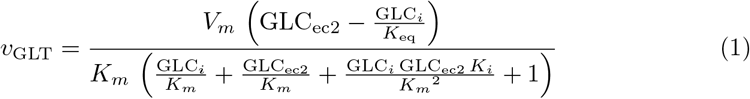

Where:

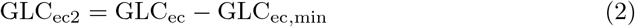

If:

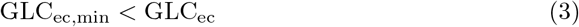

(Suarez-Mendez et al., 2014) already noticed a similar phenomenon when modelling glucose uptake dynamics. Here, we assumed that the threshold was an effect of glucose sensing under certain conditions [23]. Glucose sensing acts independently of glucose uptake [53] and is known to activate a cascade of reactions and ultimately lead to altered gene expression in yeast [54–56]. This could imply that, in line with [57], glucose sensing is observable under the FF condition, but not in the SP experiments in which residual glucose concentrations are remarkably higher [9].

### 13C-labelled metabolite mass balances validate the model but suggest caveats in carbohydrate storage metabolism

Metabolic and flux profiles agreed between simulations and experimental data (Fig 4). Even though 13C isotope labelling was used in the flux estimation [26], these data have not been implemented in kinetic models yet. Here, we aimed at validating the model by implementing individual mass balances for each labelled metabolite (carbon structures) in the network. We found a considerable degree of agreement when simulating enrichment profiles. For the first 100 s of cycle, the percentage of enriched metabolite rose to about 80% for most metabolites (Fig 7A), to then decay as recirculation of unlabelled trehalose and glycogen became more prominent (Fig 7B-D).

**Fig 7.**
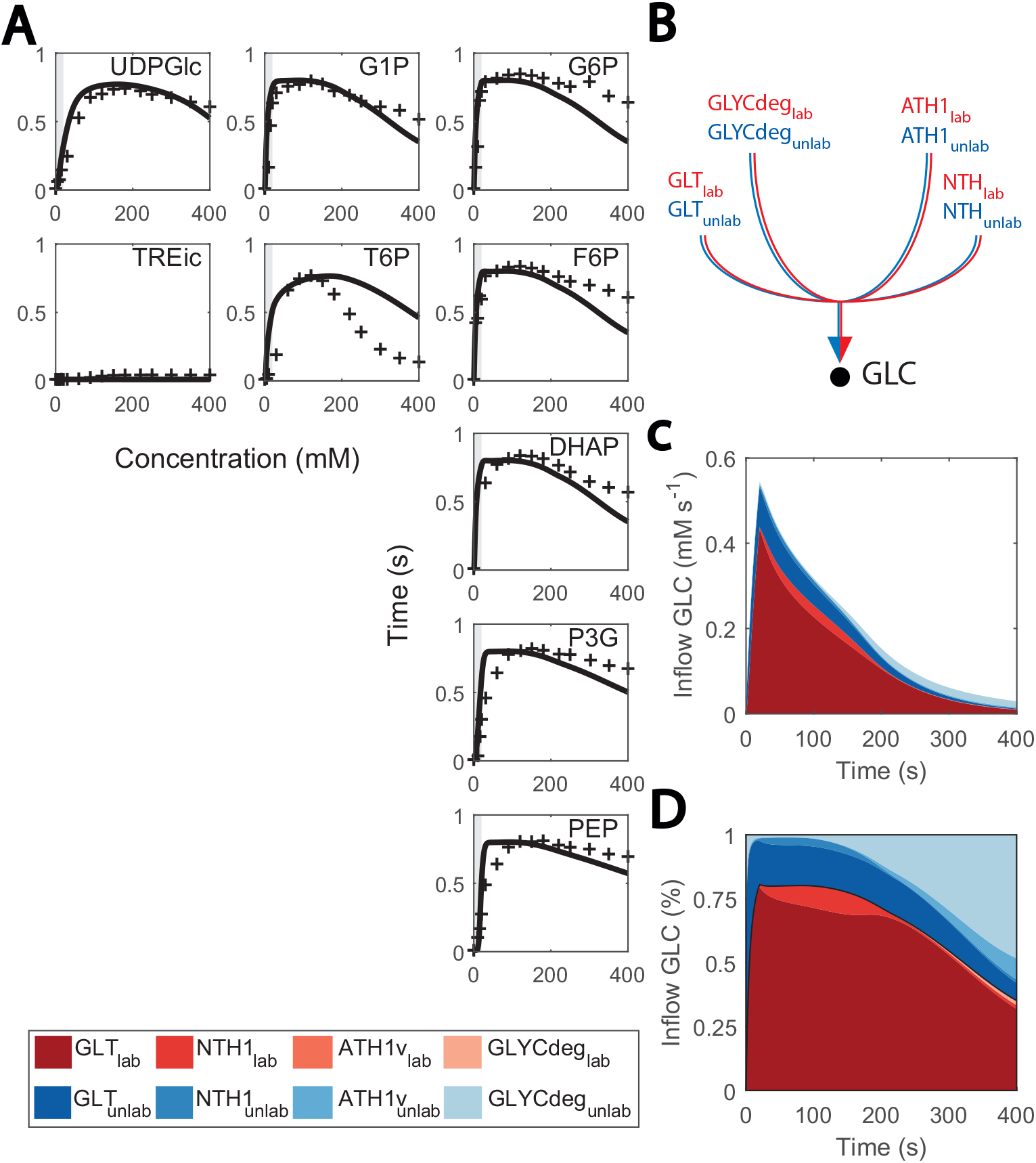
Predicted and observed 13C labelling enrichment during FF cycles. **(A) Enrichment of intracellular metabolite (%) vs time**. Black lines consist of the simulations and red markers to the experimental data points. Feast phase is shaded in grey, and famine phase has no shading. X-axis is cycle time, from 0 to 400 seconds and Y-axis is enrichment percentage, from 0 to 100%. **(B) Diagram of inflow to cytosolic glucose. (C) Fluxes that positively contribute to the cytosolic glucose mass balance (mM s**_**-1**_**) vs cycle time (s)**. Red coloured are labelled data, blue coloured, non-labelled. **(D) Contribute of each flux to the cytosolic glucose mass balance (in %) vs cycle time (s)**. Red coloured are labelled data, blue coloured, non-labelled.

Conversely, the model also showed limitations. Enrichment of T6P decreased slower and glycolytic metabolites faster than expected during the late cycle, which could indicate that there is a surplus of glycogen recirculation which was mostly unlabelled. This might be explained by the current glycogen metabolism kinetics, which were simplistic here. Small deviations from the experimental value can have a great impact on the late cycle stage, given that fluxes in the network are generally low. We initially aimed at representing glycogen synthesis and degradation as mass action or Michaelis-Menten kinetics, but unfortunately, this did not resemble the experimental reaction rates due to the high and relatively constant glycogen concentrations. Therefore, simplified phenomenological expressions were used (see details in supplementary S2 Appendix). Besides glycogen metabolism, enrichment of lower glycolysis metabolites was faster than expected, which could be attributed to the changes observed in other isoenzymes such as TDH (Fig 2).

### Identification of missing kinetic mechanisms

Another location with uncertainty is the trehalase reaction, which is carried out by an acid and neutral enzyme (ATH1 and NTH1, respectively) [58,59] and whose in vivo fluxes were quantified in the FF condition in [26]. In the model simulations, NTH1 trehalase activity was reproduced, but only if cytosolic trehalose concentration was artificially low (Fig 8A-C) and redirected to the other compartments. Nonetheless, trehalose is expected to locate more in the cytosol than the vacuole [60]. This occurred as a result of NTH1 reaction being modelled as simple Michaelis-Menten kinetics (Fig 8A) [61]. To fit these kinetics, cytosolic concentrations were kept very low with a comparatively high increase during the cycle. To further doubt the current model understanding, the estimated Km.TRE decreased from 2.11 to 0.13 mM, but it was experimentally quantified to be 3-8 mM [62]. It is not known to us what this missing regulation could be. Post translational regulation acting on NTH1 could be a possible explanation [32, 62], but new data on the state of the enzyme would be required to confirm this claim.

**Fig 8.**
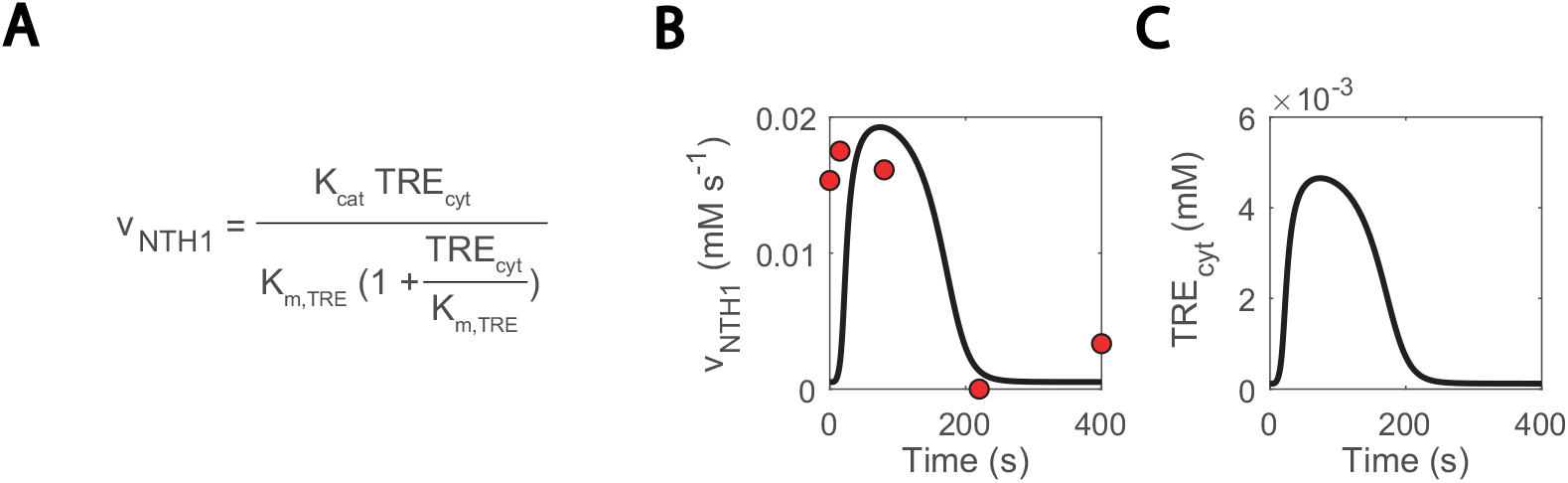
Missing regulation on NTH1 could explain excessively low simulated cytosolic trehalose concentrations. **(A) NTH1 reaction kinetics. (B) NTH1 reaction rate. (C) Cytosolic trehalose**. Blank lines show simulations and red dots experimental data.

## Conclusions and summary

In this work, we show how integrating experimental data from metabolomics, fluxomics and proteomics allowed to identify relevant mechanisms of metabolic adaptation to repetitive dynamic conditions. Especially, testing different subsets of parameters for recalibration highlighted transporters and phosphorylation reactions as crucial for the adaptation. This *in-silico* approach is comparable to the experimental approach of metabolic reverse engineering [63], but much faster and less laborious as no experiments with combinatorial genome modifications are required. The combinatorial approach can also be applied to other industrially relevant downscaling setups which are relevant to find out key parameter changes in a relatively simple manner and further optimize the bioprocess.

Furthermore, integrating labelling directly or indirectly (via flux estimation) reveals if the currently assumed mechanistic kinetics can sufficiently describe the intracellular flux and metabolome. Here, glucose uptake could not be explained by facilitated diffusion only (equation 2) but required a glucose threshold concentration (Fig 4). In addition, some reactions of the storage metabolism required non-mechanistic adjustments to reproduce the observed labelling enrichments.

## Materials and methods

### Strain and growth conditions

The haploid yeast *Saccharomyces cerevisiae* CEN PK 113-7D strain was grown at 30°C, pH5, first in a batch phase and then in chemostat at dilution rate 0.1h_-1_ [13]. The repetitive feast/famine regime began after five residence times and consisted of 20/380 seconds-cycles in which a feed was added in the first 20 seconds. The concentration of this feed was 20 times higher than the one of the chemostat phase, to ensure that the culture would overall receive the same amount of glucose. Data was collected after 20 cycles. For further reference, see [13].

### Experimental data sets used in this work

The experimental data sets used in this work consisted of metabolite concentrations measurements [13] and the respective calculated reaction rates [26]. Samples were collected more frequently during the feast than the famine phase, since network dynamics changed rapidly during the first. Extracellularly, concentrations were measured for carbohydrates glucose and trehalose. Intracellularly, concentrations were measured for carbohydrates involved in glycolysis, trehalose cycle, PPP, glycerol branch and the TCA cycle, and for adenosine nucleotides. Dynamic fluxes were estimated as piece-wise linear functions [64] using a consensus stoichiometric model for yeast [65]. Fluxes were estimated for glycolysis, carbohydrate storage metabolism (trehalose and glycogen cycles), PPP and the TCA cycle [26].

### Model description

A kinetic model of yeast central carbon metabolism was adapted in this work [29]. The original model contained the reactions that compose glycolysis, glycerol branch, a simplified trehalose cycle. Reactions of the PPP, TCA cycle and uptake of glycolytic metabolites for biomass production were lumped as sink reactions (similar to [66]). An overview of the developed model can be seen in (Fig 3). The following modifications were made to represent the complexity of carbon storage metabolism seen in the data and adapt the sink reactions of the TCA to the feast famine setup:

1. New reactions were added to represent a complete trehalose cycle and glycogen synthesis and degradation:
  - The A-glucoside transporter (AGT1) mobilizes trehalose between the extra-cellular space and cytosol [32]. Its reaction rate was modelled using reversible uni-uni MM kinetics. Since the experimental data pointed at a decay in its activity during the cycle but it did not contain any information on possible inhibitors, an inhibitory effect of T6P was added as a proxy of an increasing flux through the trehalose cycle.
  - A vacuolar transport of trehalose was added to mobilize trehalose between cytosol and vacuole-like compartments. Even though trehalose can be compartmentalized in vesicles in the cytosol, the kinetics of the process are not known. Here it was assumed that reversible MM kinetics determine this process, as with AGT1.
  - Acid trehalase (ATH1, EC 3.2.1.28) degrades trehalose to glucose. It acts in more acid environments that the cytosol such as the vacuole or the intracellular space [32], even though its location is still under debate. This reaction was modelled using irreversible MM kinetics. Similar to AGT1, inhibition by T6P was added.
  - UDP-Glucose phosphorylase (UDPG, EC 2.7.7.9) carries out the reaction from G1P to UDP-gloucose, which is later used as substrate for glycogen synthesis. This reaction was adapted from [61] and modelled using an ordered bi-bi mechanism.
  - Glycogen synthesis was not modelled by enzymatic kinetics but interpolated from the experimental data in this study, with an added UDP-glucose saturation factor.
  - Glycogen degradation was also interpolated from the experimental data in this study, with an added UDP-glucose saturation factor.
2. The sink reactions were optimized for chemostat growth [67] in the previous model. At a dilution rate of 0.1 h_-1_ the fluxes observed were higher than the ones seen under the FF regime. As a result, the flux simulated in FF towards the TCA cycle via the sink of pyruvate was over-estimated, resulting in a lesser flux towards the fermentative direction and more CO_2_ being produced than measured. A factor was added to the reaction accounting for the pyruvate sink to reduce its flux and fit the CO_2_ produced in the experiment.

In the following section, further details on model implementations are discussed.

### System of ordinary differential equations

The model consists of a series of ordinary differential equation representing the mass balances for each metabolite in the model. The model contained three compartments: cytosol, vacuole, and extracellular space. The metabolites that are part of glycolysis, trehalose and glycogen cycles, glycerol branch and cofactors metabolism were located in the cytosol, trehalose could be compartmentalized in the vacuole or secreted to the extracellular space, and glucose was modelled extracellularly as well. Meanwhile the amount of carbon structure inside the cytosol depended on the inflow of glucose and outflow of the system, moiety conservations were used for cofactors as in [22]. The sum of adenosine and nicotinamide adenine nucleotides (ATP + ADP + AMP and NAD + NADH, respectively) was kept constant in the cell. The model mass balances can be seen in detail in supplementary S1 Appendix.

### Reaction rate equations

The reaction rate equations used in this model followed Michaelis-Menten kinetics in most cases, but there were exceptions: PFK kinetics are affected by multiple regulators and the alternance between tense and relaxed state [27], PYK and PDC follow Hill-type kinetics [28] and Glucose transport occurs by facilitated diffusion (equilibrium constant equals 1). Additionally, multiple allosteric regulations occur in the network, both activation and (competitive) inhibition. Reactions were made reversible, except hydrolysis reactions, due to their remarkably negative Gibbs energy (reference) and the sink reactions in the model. Reaction rates were expressed in (mM s_-1_). The kinetic rate expressions can be seen in detail in supplementary S2 Appendix and a summary on each of them in supplementary S4 Appendix.

### Simulation setup

The simulations were performed in three steps aimed at resembling the experimental process that cells underwent in the experiments in [13]. The first step consisted of a chemostat, which also served to confirm that the system remained in a physiological realistic steady state. The second step consisted of the feast famine cycles. 20 repetitive cycles were simulated in which glucose was fed for the first 20 seconds of cycle without any outgoing flux. For the rest of the cycle, no glucose was fed, and the outgoing flux lasted until the same amount of volume increased in the first 20 seconds had been emptied, by approximately 260 seconds of cycle. Afterwards, both incoming and outgoing fluxes were kept at zero. After running for 20 cycles, the resulting simulation was compared to the experimental metabolite concentrations in reaction rates obtained in [13]. The third step concerned the simulation of the enrichment profiles reported in [26]. This extra simulation was run after the second step and 99% of the inflow of glucose was ^13^C labelled. All the simulations were carried with the abovementioned mechanistic model. Matlab version 9.3.0.713579, R2017b, and the ode15s solver were used. A graphical view of how simulations were constructed can be found in supplementary S5 Appendix.

### Implementation of 13C-labeling data simulations

As described by [25], reactions of a metabolic network can be correctly represented with mass isotopomeric models if there are no cleavage reactions present, because in the latter the position of the labelled carbon(s) is decisive to define the isotopomers of the following metabolites. In the interest of providing an accurate model without over-complicating it, the kinetic model is thus expanded to labelled carbon enrichment, instead of simulations of isotopic transients, as the simulation of the isotopic transients, requires determination of the distribution of the possible C-labelled atoms for every metabolite of the network and additionally accounting for the bidirectionally of reactions in the isotopomer balance equations, which is no longer as trivial as building an enrichment model [68].

For each carbon-based metabolite in the model, a mass balance was added to account for its respective ^13^C-labelled fraction. In this secondary mass balance, the input and output reaction rates were the same than for the total metabolite concentration mass balance, but it was multiplied by the respective fraction of labelled metabolite. Whereas the metabolite total concentrations depend solely on the enzymatic rates, the metabolite labelled concentrations also depend on the fractions of other labelled metabolites (labelled concentration / total concentration) that the reactions use as substrate. Moreover, now the enzymatic fluxes need to be adjusted for reversible enzymatic reactions.

For mass balances equations of total metabolite concentrations, reversible enzymatic fluxes are defined with a positive value in one direction and negative in the reverse direction. However, to implement the mass balances equations of labelled metabolite concentrations, these are multiplied by metabolite’s labelled fractions and so they need to be always positive. Thus, enzymatic fluxes that change directions during the simulation, are implemented as a forward and backwards flux. These are only used in the equations for labelled metabolite concentrations and are defined in the model with conditional statements. An example of the mass balance of labelled and unlabelled acetate can be seen below. Reaction reversibility is already accounted for inside the calculation of the reaction rate:

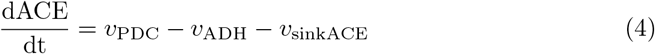

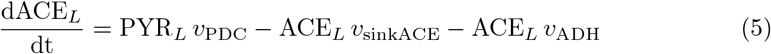

Where L refers to the labelled metabolite fraction. Enrichment simulations were performed after the 20 repetitive cycles were simulated. The experimental data consisted of percentage level of ^13^C enrichment over cycle time, obtained from [26].

### Parameter values used in this work

The initial parameter values were obtained from the original model [29]. These parameter values had been fitted to experimental data (metabolomics and fluxomics) from a data at different steady states [67] and SRE [9]. A subset of the parameter values including enzymes HXT and GLK, the ones involved in the trehalose cycle and ATPase kinetics were estimated in this work. The values of the kinetic rate expressions used in this work can be seen in supplementary S3 Appendix.

### Estimation of in vivo parameters

Some reactions in the model underwent changes during the feast famine cycles. For instance, the proportions of the isoenzymes HXK/GLK changed. To account for its effect on kinetic parameters such as *K*_*M*_ or *K*_*cat*_, reaction parameters were estimated for HXT and HXK/GLK. Additionally, trehalose cycles parameters were estimated, since the final structure of the cycle was different that the initial, and the ATPase reaction rate constant needed to change as well. For initial parameter guesses, the initial parameter values from [29] were used. The nonlinear least-squares solver lsqnonlin from the Optimization Toolbox, using an interior reflective Newton method [69], was used to estimate the parameters by minimizing the error between measured and simulated data during the transient experiment.

### Design of the cost functions: combination of enzymes and weighting factors

Experimental quantification of isoenzymes pointed at the couple HXK/GLK experiencing the biggest deviation prior and after the FF cycles, but minor changes were also confirmed for the other enzymes in glycolysis (Figure 2). Still, estimation of all the kinetic constants in glycolysis simultaneously was undesirable due to the appearance of parameter dependencies [48] which could lead to unphysiological parameter values. Therefore, to describe the changes in the experimental data with only the essential number of enzymes changing, the parameters were fitted to the data in multiple occasions. In each of those, a different selection of enzymes and cost function weighting factors was used:

- Selection of enzymes: multiple enzyme combinations were tested. These combinations contained the trehalose cycle and added different enzymes from glycolysis each time. The combination selected was the one that described experimental data properly while making physiological sense (such as including the changes in HXK/GLK) and having the smallest number of enzymes possible. Simultaneously, random combinations of enzymes were also tested, to confirm results and give robustness to the method.
- Combination of weighting factors: it was not clear at first if it would be possible to describe all the experimental data simultaneously. For this purpose, each of the abovementioned enzyme combinations was run repeated times, each of them with a different set of weighting factors. The errors for every metabolite were normalized so that they would contribute with the same weight to the cost function. Additional weighting factors changed these weights in three orders of magnitude at most.

### Design of the cost functions: regularization

Parameter dependencies could still appear for a selection of enzymes or within a single enzyme, even though less generalized than if all enzymes had been optimized together. To avoid this problem, parameters were estimated again after the previous round of data fit which was only based on selecting enzymes and cost function weights. This time L1-type regularization was implemented to force the parameter estimates closer to the initial parameter set, as long as experimental data could be properly fit, helping to identify the important parameter changes for a specific dataset [36, 37]. In this way, only the necessary parameters needed to change to fit the FF data were made distinguishable. The regularization factor λ was applied to the cost function in the following manner:

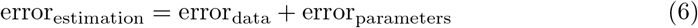

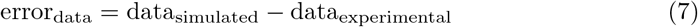

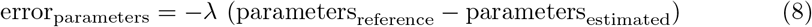

## Supporting information

**S1 Fig. Simulation of metabolic concentrations is robust to parameter changes within 10% of their estimated value**. 2D histogram plot of 10.000 model samples. Darker areas point at more dense regions. Models were sampled by adding random noise to all the parameters in the network, within a range of ±10% of the original parameter value.

**S2 Fig. Simulation of reaction rates is robust to parameter changes within 10% of their estimated value**. 2D histogram plot of 10.000 model samples. Darker areas point at more dense regions. Models were sampled by adding random noise to all the parameters in the network, within a range of ±10% of the original parameter value.

**S3 Fig. Metabolite concentrations (mM) over time (s). Model fit**. (Gray) Non-regularized, (black) regularized and (black dots) experimental data. Intracellular trehalose was experimentally measured and a 9/1-distribution ratio was assumed for the vacuolar and cytosolic concentrations.

**S4 Fig. Reaction rates (mM s**^**-1**^**) over time (s). Model fit**. (Gray) Non-regularized, (black) regularized and (black dots) experimental data.

**S5 Fig. Glucose sensing is needed to explain HXT kinetics: Individual parameter effect**. Glucose uptake rate at 400 vs at 20 second. Blue and yellow colours show smaller and bigger parameter values. 1000 samples were run, within 3 orders of magnitude above and below the estimated parameters. Parameters were randomized for HXT kinetics and external glucose concentration was fit to the experimental data. The red dot consists in the experimental data point.

**S1 Appendix. Ordinary differential equations**.

**S2 Appendix. Rate equations**.

**S3 Appendix. Parameter set**.

**S4 Appendix. Summary rate equations**.

**S5 Appendix. Simulation setup**.

## Data availability statement

The data used in this work can be found in the github repository of the author (github.com/DavidLaoM/y3m2_ff). The experimental proteomics dataset can be found in [39].

## Author Contributions

- **Conceptualization:** David Lao-Martil, Koen J.A. Verhagen, S. Aljoscha Wahl.
- **Formal analysis:** David Lao-Martil, Koen J.A. Verhagen.
- **Funding acquisition:** Natal A. W. van Riel, S. Aljoscha Wahl.
- **Investigation:** David Lao-Martil, Koen J.A. Verhagen, Ana H. Valdeira Caetano, Ilse Pardijs.
- **Methodology:** David Lao-Martil, Koen J.A. Verhagen, Ana H. Valdeira Caetano, Ilse Pardijs.
- **Software:** David Lao-Martil, Koen J.A. Verhagen, Ana H. Valdeira Caetano, Ilse Pardijs.
- **Supervision:** Natal A. W. van Riel, S. Aljoscha Wahl.
- **Visualization:** David Lao-Martil, Koen J.A. Verhagen.
- **Writing - original draft:** David Lao-Martil, Koen J.A. Verhagen.
- **Writing - review & editing:** David Lao-Martil, Koen J.A. Verhagen, Natal A.W. van Riel, S. Aljoscha Wahl.

## Acknowledgements

The authors would like to acknowledge Dr Joep Schmitz for assistance with sample processing.

