## supplementary_materials for "Using kinetic modelling to infer adaptations in *Saccharomyces cerevisiae* carbohydrate storage metabolism to feast famine regimes": S2_Appendix.pdf

### Rate Equations

The following kinetic expressions were used:

$$v_{\mathrm{GLT}} = \frac{V_m \left(\mathrm{GLCo} - \frac{\mathrm{GLCi}}{K_{\mathrm{eq}}}\right)}{K_m \left(\frac{\mathrm{GLCi}}{K_m} + \frac{\mathrm{GLCo}}{K_m} + \frac{\mathrm{GLCi}\,\mathrm{GLCo}\,K_i}{{K_m}^2} + 1\right)} \tag{1}$$

$$v_{\mathrm{HXK}} = \frac{V_m \left(\mathrm{ATP}\,\mathrm{GLCi} - \frac{\mathrm{ADP}\,\mathrm{G6P}}{K_{\mathrm{eq}}}\right)}{K_{m,\mathrm{glc}}\,K_{m,\mathrm{atp}} \left(\frac{\mathrm{ADP}}{K_{m,\mathrm{adp}}} + \frac{\mathrm{ATP}}{K_{m,\mathrm{atp}}} + 1\right) \left(\frac{\mathrm{G6P}}{K_{m,\mathrm{g6p}}} + \frac{\mathrm{GLCi}}{K_{m,\mathrm{glc}}} + \frac{\mathrm{T6P}}{K_{i,\mathrm{t6p}}} + 1\right)} \tag{2}$$

$$v_{\mathrm{PGM1}} = \frac{V_m \left(\mathrm{G1P} - \frac{\mathrm{G6P}}{K_{\mathrm{eq}}}\right)}{K_{m,\mathrm{G1P}} \left(\frac{\mathrm{G1P}}{K_{m,\mathrm{G1P}}} + \frac{\mathrm{G6P}}{K_{m,\mathrm{G6P}}} + 1\right)} \tag{3}$$

$$v_{\mathrm{TPS1}} = \frac{V_m\,\mathrm{F6P}\,\mathrm{G6P}\,\mathrm{UDPGLC}}{K_{m,\mathrm{G6P}}\,K_{m,\mathrm{UDPGLC}} \left(\frac{\mathrm{G6P}}{K_{m,\mathrm{G6P}}} + 1\right) \left(\frac{\mathrm{PI}}{K_{i,\mathrm{PI}}} + 1\right) \left(\frac{\mathrm{UDPGLC}}{K_{m,\mathrm{UDPGLC}}} + 1\right) (\mathrm{F6P} + K_{m,\mathrm{F6P}})} \tag{4}$$

$$v_{\mathrm{TPS2}} = \frac{V_m\,\mathrm{PI}\,\mathrm{T6P}}{\mathrm{PI}\,(K_{m,\mathrm{T6P}} + \mathrm{T6P}) + K_{i,\mathrm{PI}}\,K_{m,\mathrm{T6P}}} \tag{5}$$

$$v_{\mathrm{NTH1}} = \frac{V_m\,\mathrm{TRE}}{K_{m,\mathrm{TRE}} \left(\frac{\mathrm{TRE}}{K_{m,\mathrm{TRE}}} + 1\right)} \tag{6}$$

$$v_{\mathrm{PGI}} = \frac{V_m \left(\mathrm{G6P} - \frac{\mathrm{F6P}}{K_{\mathrm{eq}}}\right)}{K_{m,\mathrm{g6p}} \left(\frac{\mathrm{F6P}}{K_{m,\mathrm{f6p}}} + \frac{\mathrm{G6P}}{K_{m,\mathrm{g6p}}} + 1\right)} \tag{7}$$

$$v_{\mathrm{PFK}} = \frac{V_m\,R\,g_R\,\mathrm{lam}_1\,\mathrm{lam}_2}{R^2 + L\,T^2} \tag{8}$$

$$\mathrm{lam}_1 = \frac{\mathrm{F6P}}{K_{R,\mathrm{F6P}}} \tag{9}$$

$$\mathrm{lam}_2 = \frac{\mathrm{ATP}}{K_{R,\mathrm{ATP}}} \tag{10}$$

$$R = \mathrm{lam}_1\,\mathrm{lam}_2 + g_R\,\mathrm{lam}_1\,\mathrm{lam}_2 + 1 \tag{11}$$

$$T = c_{\mathrm{ATP}}\,\mathrm{lam}_2 + 1 \tag{12}$$

$$L = \frac{L_0 \left(\frac{\mathrm{AMP}\,c_{i,\mathrm{AMP}}}{K_{\mathrm{AMP}}} + 1\right)^2 \left(\frac{\mathrm{ATP}\,c_{i,\mathrm{ATP}}}{K_{\mathrm{ATP}}} + 1\right)^2 \left(\frac{\mathrm{F26bP}\,c_{i,\mathrm{F26bp}}}{K_{\mathrm{F26bP}}} + \frac{\mathrm{FBP}\,c_{i,\mathrm{FBP}}}{K_{\mathrm{FBP}}} + 1\right)}{\left(\frac{\mathrm{AMP}}{K_{\mathrm{AMP}}} + 1\right)^2 \left(\frac{\mathrm{ATP}}{K_{\mathrm{ATP}}} + 1\right)^2 \left(\frac{\mathrm{F26bP}}{K_{\mathrm{F26bP}}} + \frac{\mathrm{FBP}}{K_{\mathrm{FBP}}} + 1\right)} \tag{13}$$

$$v_{\mathrm{ALD}} = \frac{V_m \left(\mathrm{FBP} - \frac{\mathrm{DHAP}\,\mathrm{GAP}}{K_{\mathrm{eq}}}\right)}{K_{m,\mathrm{FBP}} \left(\frac{\mathrm{FBP}}{K_{m,\mathrm{FBP}}} + \left(\frac{\mathrm{DHAP}}{K_{m,\mathrm{dhap}}} + 1\right) \left(\frac{\mathrm{GAP}}{K_{m,\mathrm{gap}}} + 1\right)\right)} \tag{14}$$

$$v_{\mathrm{TPI}} = \frac{V_m \left(\mathrm{DHAP} - \frac{\mathrm{GAP}}{K_{\mathrm{eq}}}\right)}{K_{m,\mathrm{dhap}} \left(\frac{\mathrm{DHAP}}{K_{m,\mathrm{dhap}}} + \frac{\mathrm{GAP}}{K_{m,\mathrm{gap}}} + 1\right)} \tag{15}$$

$$v_{\mathrm{GPD}} = \frac{V_m \left(\mathrm{DHAP}\,\mathrm{NADH} - \frac{\mathrm{G3P}\,\mathrm{NAD}}{K_{\mathrm{eq}}}\right) \text{[illegible]}}{K_{m,\mathrm{NADH}}\, K_{m,\mathrm{DHAP}} \left(\frac{\mathrm{NAD}}{K_{m,\mathrm{NAD}}} + \frac{\mathrm{NADH}}{K_{m,\mathrm{NADH}}} + 1\right) \left(\frac{\mathrm{DHAP}}{K_{m,\mathrm{DHAP}}} + \frac{\mathrm{G3P}}{K_{m,\mathrm{G3P}}} + 1\right) \left(\frac{\mathrm{ADP}}{K_{i,\mathrm{ADP}}} + \text{[illegible]}\right.} \tag{16}$$

$$v_{\mathrm{HOR2}} = \frac{V_m\, \mathrm{G3P}}{K_{m,\mathrm{G3P}} \left(\frac{\mathrm{G3P}}{K_{m,\mathrm{G3P}}} + 1\right) \left(\frac{\mathrm{PI}}{K_{i,\mathrm{PI}}} + 1\right)} \tag{17}$$

$$V_{\mathrm{GLYCt}} = K_{\mathrm{GLYCt}} \left(\mathrm{GLYC} - \mathrm{GLYCe}\right) \tag{18}$$

$$v_{\mathrm{GAPDH}} = \frac{V_m \left(\mathrm{GAP}\,\mathrm{NAD}\,\mathrm{PI} - \frac{\mathrm{BPG}\,\mathrm{NADH}}{K_{\mathrm{eq}}}\right) \text{[illegible]}}{K_{m,\pi}\, K_{m,\mathrm{nad}}\, K_{m,\mathrm{gap}} \left(\left(\frac{\mathrm{BPG}}{K_{m,\mathrm{bpg}}} + 1\right) \left(\frac{\mathrm{NADH}}{K_{m,\mathrm{nadh}}} + 1\right) + \left(\frac{\mathrm{GAP}}{K_{m,\mathrm{gap}}} + 1\right) \left(\frac{\mathrm{NAD}}{K_{m,\mathrm{nad}}} + 1\right) \text{[illegible]}\right.} \tag{19}$$

$$v_{\mathrm{PGK}} = \frac{V_m \left(\mathrm{ADP}\,\mathrm{BPG}\,K_{\mathrm{eq}} - \mathrm{ATP}\,\mathrm{P3G}\right)}{K_{m,\mathrm{P3G}}\, K_{m,\mathrm{ATP}} \left(\frac{\mathrm{BPG}}{K_{m,\mathrm{BPG}}} + \frac{\mathrm{P3G}}{K_{m,\mathrm{P3G}}} + 1\right) \left(\frac{\mathrm{ADP}}{K_{m,\mathrm{ADP}}} + \frac{\mathrm{ATP}}{K_{m,\mathrm{ATP}}} + 1\right)} \tag{20}$$

$$v_{\mathrm{PGM}} = \frac{V_m \left(\mathrm{P3G} - \frac{\mathrm{P2G}}{K_{\mathrm{eq}}}\right)}{K_{m,\mathrm{P3G}} \left(\frac{\mathrm{P2G}}{K_{m,\mathrm{P2G}}} + \frac{\mathrm{P3G}}{K_{m,\mathrm{P3G}}} + 1\right)} \tag{21}$$

$$v_{\mathrm{ENO}} = \frac{V_m \left(\mathrm{P2G} - \frac{\mathrm{PEP}}{K_{\mathrm{eq}}}\right)}{K_{m,\mathrm{P2G}} \left(\frac{\mathrm{P2G}}{K_{m,\mathrm{P2G}}} + \frac{\mathrm{PEP}}{K_{m,\mathrm{PEP}}} + 1\right)} \tag{22}$$

$$v_{\mathrm{PYK}} = \frac{V_m\, \mathrm{ADP}\,\mathrm{PEP} \left(\frac{\mathrm{PEP}}{K_{m,\mathrm{pep}}} + 1\right)^{\mathrm{hill}}}{K_{m,\mathrm{adp}}\, K_{m,\mathrm{pep}} \left(\left(\frac{\mathrm{PEP}}{K_{m,\mathrm{pep}}} + 1\right)^{\mathrm{hill}} + L \left(\frac{\frac{\mathrm{ATP}}{K_{i,\mathrm{ATP}}} + 1}{\frac{\mathrm{FBP}}{K_{a,\mathrm{FBP}}} + 1}\right)^{\mathrm{hill}}\right) \left(\frac{\mathrm{ADP}}{K_{m,\mathrm{adp}}} + 1\right) \left(\frac{\mathrm{PEP}}{K_{m,\mathrm{pep}}} + 1\right)} \tag{23}$$

$$v_{\mathrm{PDC}} = \frac{V_m \left(\frac{\mathrm{PYR}}{K_{m,\mathrm{PYR}}}\right)^{\mathrm{hill}}}{\left(\frac{\mathrm{PYR}}{K_{m,\mathrm{PYR}}}\right)^{\mathrm{hill}} + 1} \tag{24}$$

$$v_{\mathrm{ADH}} = -\frac{\text{[illegible]}}{K_{i,\mathrm{NAD}}\, K_{m,\mathrm{ETOH}} \left(\frac{\mathrm{NAD}}{K_{i,\mathrm{NAD}}} + \frac{\mathrm{NADH}}{K_{i,\mathrm{NADH}}} + \frac{\mathrm{ACE}\, K_{m,\mathrm{NADH}}}{K_{i,\mathrm{NADH}}\, K_{m,\mathrm{ACE}}} + \frac{\mathrm{ACE}\,\mathrm{NADH}}{K_{i,\mathrm{NADH}}\, K_{m,\mathrm{ACE}}} + \frac{\mathrm{ETOH}\, K_m \text{[illegible]}}{K_{i,\mathrm{NAD}}\, K_m \text{[illegible]}} \text{[illegible]}\right.} \tag{25}$$

$$V_{\mathrm{ETOHt}} = K_{\mathrm{ETOHt}} \left(\mathrm{ETOH} - \mathrm{ETOHe}\right) \tag{26}$$

$$V_{\text{mito}} = \frac{V_m \,\text{ADP}\,\text{PI}}{(\text{ADP} + K_{m,\text{ADP}})\,(K_{m,\text{PI}} + \text{PI})} \tag{27}$$

$$V_{\text{ATPase}} = \frac{\text{ATP}\,K}{\text{ADP}} \tag{28}$$

$$V_{\text{ADK}} = K\left(\text{ADP}^2 - \frac{\text{AMP}\,\text{ATP}}{K_{\text{eq}}}\right) \tag{29}$$

$$V_{\text{vacPi}} = -K\,(\text{PI} - \text{PIvac}) \tag{30}$$

$$V_{\text{mitoNADH}} = \frac{V_m \,\text{NADH}}{K_m + \text{NADH}} \tag{31}$$

$$V_{\text{sinkG6P}} = \frac{V_m \,\text{G6P}}{\text{G6P} + K_m} \tag{32}$$

$$V_{\text{sinkF6P}} = \frac{V_m \,\text{F6P}}{\text{F6P} + K_m} \tag{33}$$

$$V_{\text{sinkGAP}} = \frac{V_m \,\text{GAP}}{\text{GAP} + K_m} \tag{34}$$

$$V_{\text{sinkP3G}} = \frac{V_m \,\text{P3G}}{K_m + \text{P3G}} \tag{35}$$

$$V_{\text{sinkPEP}} = \frac{V_m \,\text{PEP}}{K_m + \text{PEP}} \tag{36}$$

$$V_{\text{sinkPYR}} = \frac{V_m \,\text{PYR}}{K_m + \text{PYR}} \tag{37}$$

$$V_{\text{sinkACE}} = \frac{V_m \,\text{ACE}}{\text{ACE} + K_m} \tag{38}$$

Reactions added in this model:

$$v_{\text{ATH}} = \frac{\text{ATH}\,K_{\text{cat}}\,\text{TRE}}{K_{M,\text{TRE}}\left(\frac{\text{TRE}}{K_{M,\text{TRE}}} + 1\right)} \tag{39}$$

$$v_{\text{AGT1}} = \frac{\text{AGT}_1\,K_{\text{cat}}\left(\text{TRE}_{\text{cyt}} - \frac{\text{TRE}_{\text{ec}}}{K_{\text{eq}}}\right)}{K_{M,\text{TRE}}\left(\frac{\text{TRE}_{\text{ec}}}{K_{M,\text{TRE}}} + \frac{\text{TRE}_{\text{cyt}}}{K_{M,\text{TRE}}} + \frac{\text{UDP}_{\text{GLC}}}{K_{i,\text{UDPglc}}} + 1\right)} \tag{40}$$

$$v_{\text{vacuoleT}} = \frac{V_{\max}\left(\text{TRE}_{\text{cyt}} - \frac{\text{TRE}_{\text{vac}}}{K_{\text{eq}}}\right)}{K_{M,\text{TRE}}\left(\frac{\text{TRE}_{\text{vac}}}{K_{M,\text{TRE}}} + \frac{\text{TRE}_{\text{cyt}}}{K_{M,\text{TRE}}} + 1\right)} \tag{41}$$

Glycogen synthesis and degradation were directly interpolated from the data, with the addition of a saturation component, to avoid that it become a constraint in the system, especially when fluxes were overall small.

$$v_{\text{glyc,synthesis}} = \frac{v_{\text{glyc,synthesis,interpolated}}\,\text{UDP}_{\text{GLC}}}{\text{UDP}_{\text{GLC}} + 1\text{E}^{-4}} \tag{42}$$

$$v_{\text{glyc,degradation}} = \frac{v_{\text{glyc,degradation,interpolated}}\,\text{Glycogen}}{\text{Glycogen} + 1\text{E}^{-4}} \tag{43}$$
