## supplementary_materials for "Using kinetic modelling to infer adaptations in *Saccharomyces cerevisiae* carbohydrate storage metabolism to feast famine regimes": S2_Fig.pdf

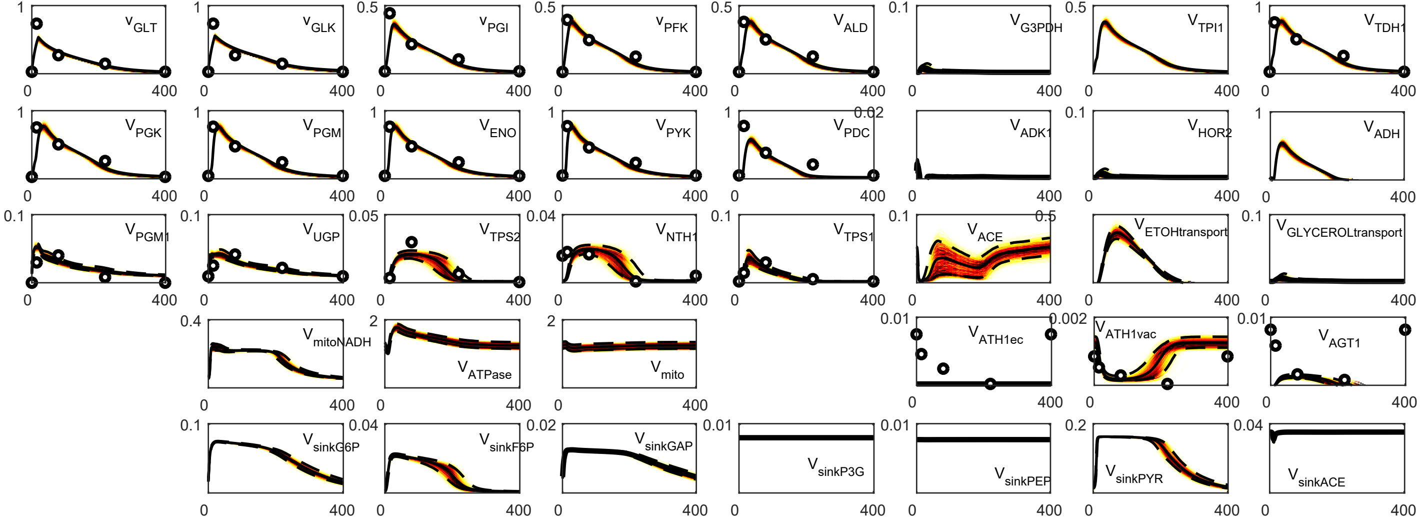


**Figure S2. Simulation of reaction rates is robust to parameter changes within 10% of their estimated value.**
2D histogram plot of 10.000 model samples. Darker areas point at more dense regions. Models were sampled by adding random noise to all the parameters in the network, within a range of ±10% of the original parameter value.
