## supplementary_materials for "Using kinetic modelling to infer adaptations in *Saccharomyces cerevisiae* carbohydrate storage metabolism to feast famine regimes": S3_Fig.pdf

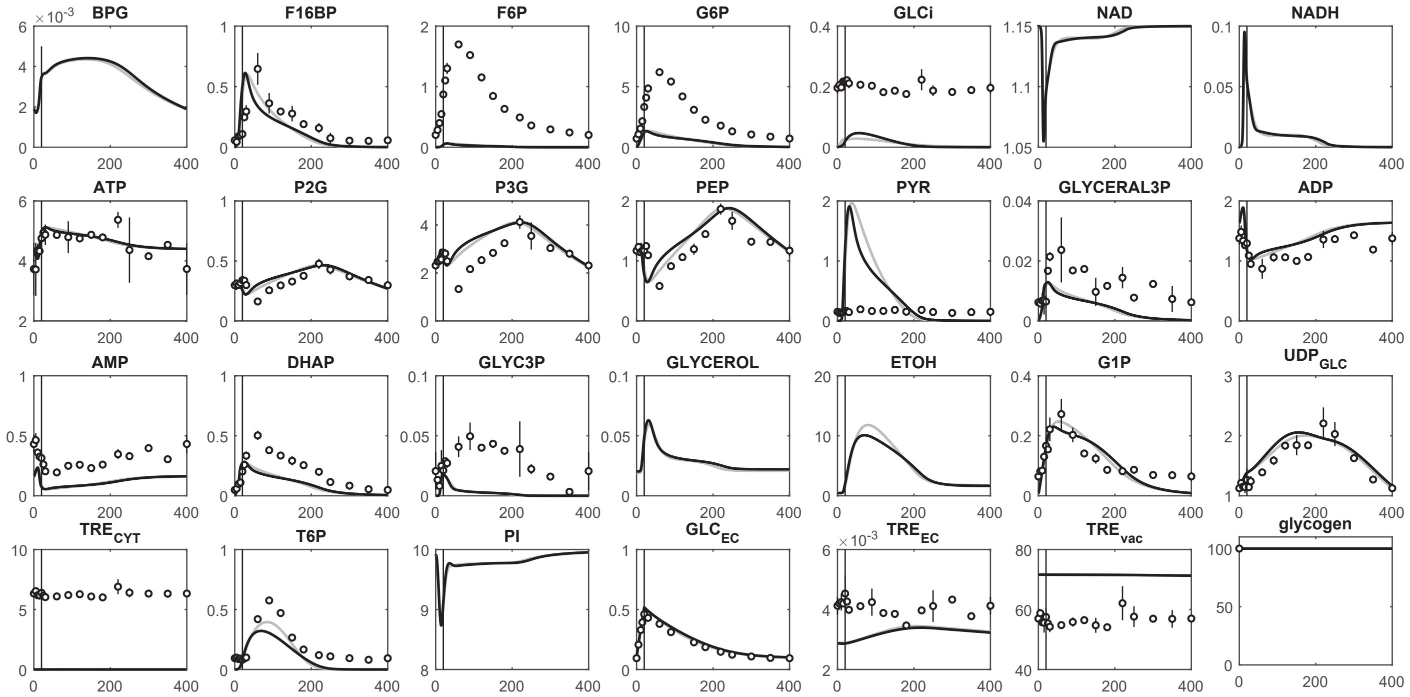


**Figure S3. Metabolite concentrations (mM) over time (s). Model fit.**
(Gray) Non-regularized, (black) regularized and (black dots) experimental data.
