## supplementary_materials for "Using kinetic modelling to infer adaptations in *Saccharomyces cerevisiae* carbohydrate storage metabolism to feast famine regimes": S4_Fig.pdf

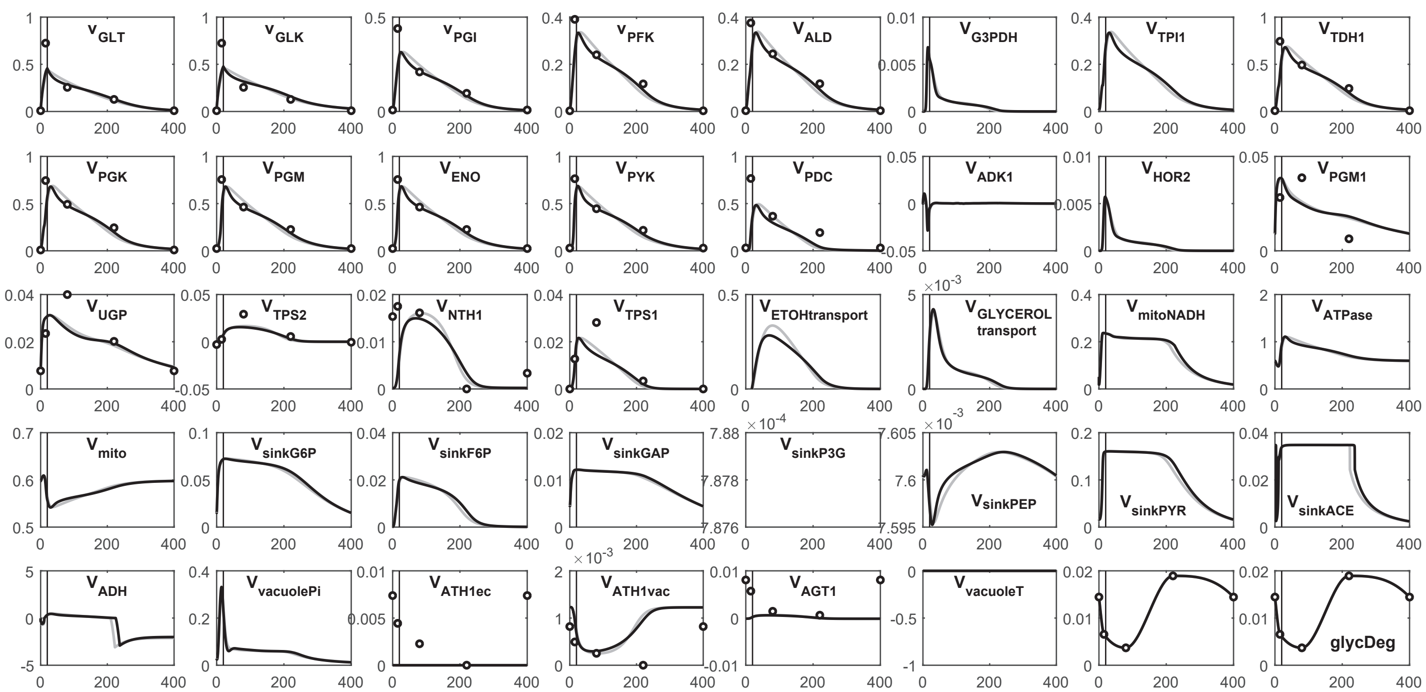


**Figure S4. Reaction rates (mM s$^{-1}$) over time (s). Model fit.**

(Gray) Non-regularized, (black) regularized and (black dots) experimental data.
