## supplementary_materials for "Using kinetic modelling to infer adaptations in *Saccharomyces cerevisiae* carbohydrate storage metabolism to feast famine regimes": S5_Appendix.pdf

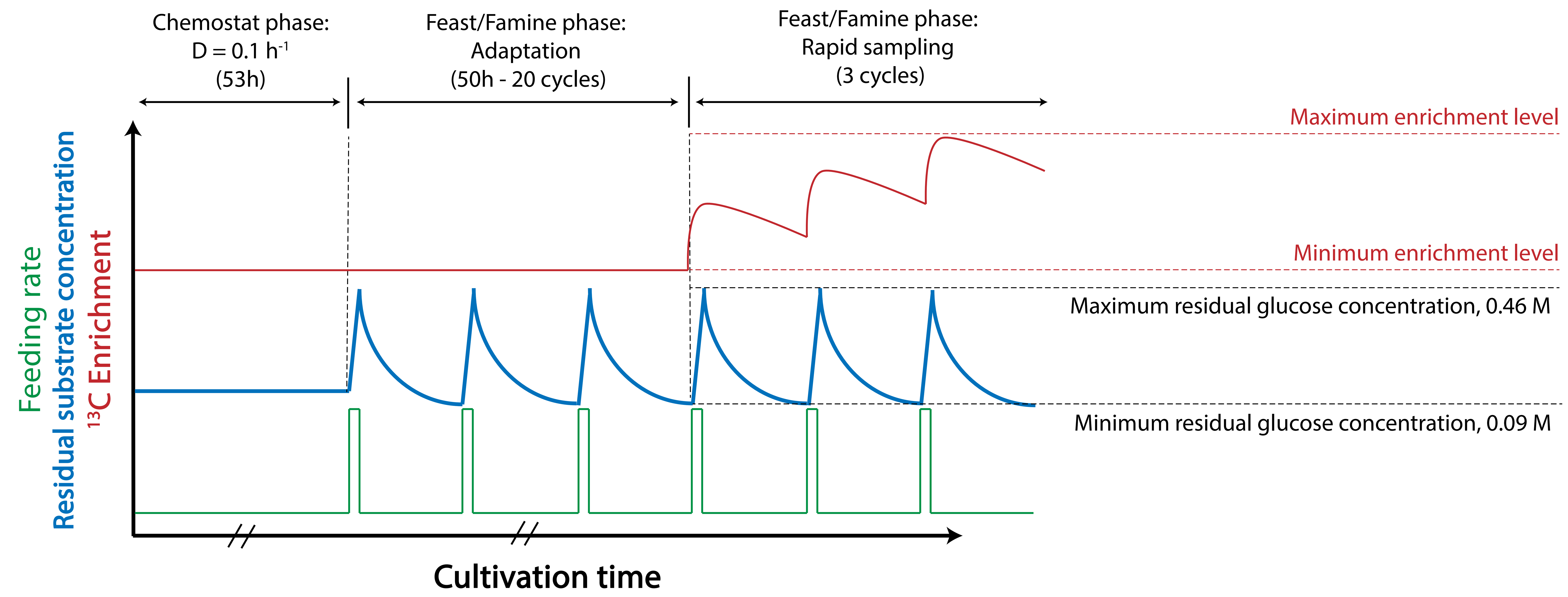


Chemostat phase

$Fin = 1.0817 \cdot 10^{-4}$ (L s$^{-1}$)

Fin = Fout

$Cx = 3.639$ ($g_{DW}$ L$^{-1}$)

Feeding phase (0-20s)

$Fin = 2.2 \cdot 10^{-3}$, Fout = 0 (L s$^{-1}$)

No feeding phase (20-280s)

Fin = 0, $Fout = 1.7 \cdot 10^{-4}$ (L s$^{-1}$)

No feeding phase (280-400s)

Fin = 0, Fout = 0 (L s$^{-1}$)

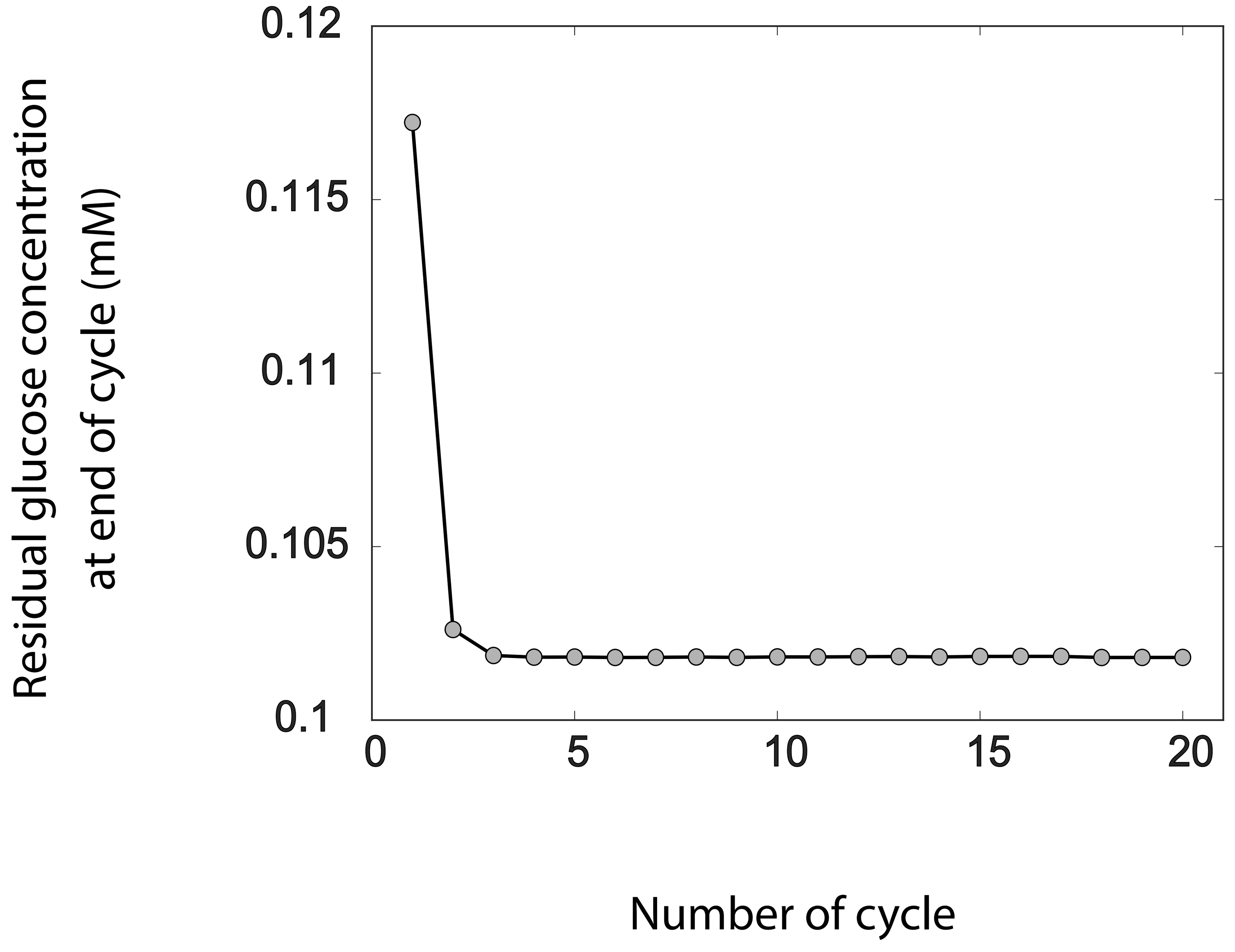
