## supplementary_materials for "Using kinetic modelling to infer adaptations in *Saccharomyces cerevisiae* carbohydrate storage metabolism to feast famine regimes": S5_Fig.pdf

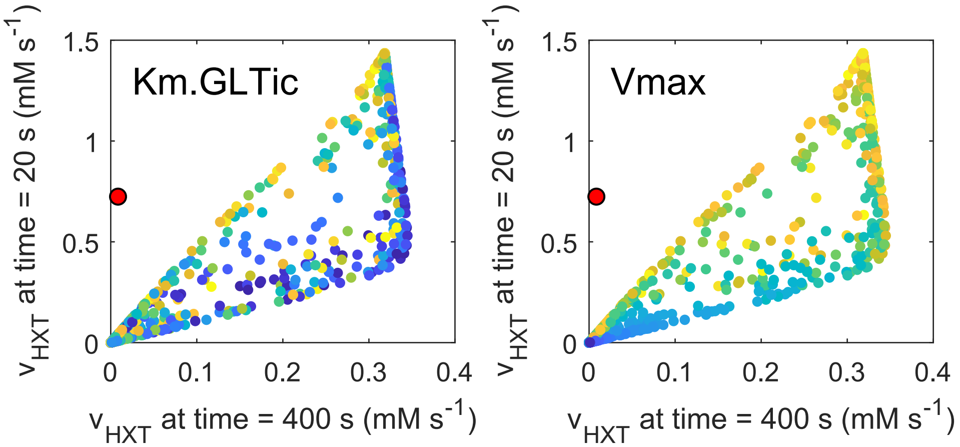


**Figure S5. Glucose sensing is needed to explain HXT kinetics: Individual parameter effect.**
Glucose uptake rate at 400 vs at 20 second. Blue and yellow colours show smaller and bigger parameter values. 1000 samples were run, within 3 orders of magnitude above and below the estimated parameters. Parameters were randomized for HXT kinetics and external glucose concentration was fit to the experimental data. The red dot consists in the experimental data point.
